# LEVIATHAN: efficient discovery of large structural variants by leveraging long-range information from Linked-Reads data

**DOI:** 10.1101/2021.03.25.437002

**Authors:** Pierre Morisse, Fabrice Legeai, Claire Lemaitre

## Abstract

Linked-Reads technologies, popularized by 10x Genomics, combine the high-quality and low cost of short-reads sequencing with a long-range information by adding barcodes that tag reads originating from the same long DNA fragment. Thanks to their high-quality and long-range information, such reads are thus particularly useful for various applications such as genome scaffolding and structural variant calling. As a result, multiple structural variant calling methods were developed within the last few years. However, these methods were mainly tested on human data, and do not run well on non-human organisms, for which reference genomes are highly fragmented, or sequencing data display high levels of heterozygosity. Moreover, even on human data, most tools still require large amounts of computing resources. We present LEVIATHAN, a new structural variant calling tool that aims to address these issues, and especially better scale and apply to a wide variety of organisms. Our method relies on a barcode index, that allows to quickly compare the similarity of all possible pairs of regions in terms of amount of common barcodes. Region pairs sharing a sufficient number of barcodes are then considered as potential structural variants, and complementary, classical short reads methods are applied to further refine the breakpoint coordinates. Our experiments on simulated data underline that our method compares well to the state-of-the-art, both in terms of recall and precision, and also in terms of resource consumption. Moreover, LEVIATHAN was successfully applied to a real dataset from a non-model organism, while all other tools either failed to run or required unreasonable amounts of resources. LEVIATHAN is implemented in C++, supported on Linux platforms, and available under AGPL-3.0 License at https://github.com/morispi/LEVIATHAN.

## 1 Introduction

Structural variants (SVs) represent variations in the structure of an organism’s genome. Detecting such events is crucial, since many of them are associated with genetic diseases. Classical short-read SV calling methods usually rely on their alignment against a reference genome, and on the detection of discordant paired-read or split read signals, in order to determine the breakpoints and types of the SVs. However, due to the limited size of the short-reads, many SVs remain undetected, while many False Positive calls are reported by such methods.

Linked-Reads technologies rely on partitioning and barcoding of diluted high-molecular-weight DNA using a microfluidic device prior to classical short-read sequencing. Molecule sizes usually range between 10 and 50 kbp on average. However, the short-reads coverage of each molecule is usually low. Indeed, for a typical 30x sequencing depth experiment, the coverage of the reference genome by the large molecules is of about 150x, but each molecule displays a weak short-read coverage of about 0.2x. 10x Genomics popularized this technology [1], but since discontinued the sales of their Linked-Reads product lines. However, large volumes of data were produced and still need to be properly analyzed, and other technologies such as TELL-Seq [2] and Haplotagging [3] emerged, and also allow the sequencing of Linked-Reads.

Thanks to the barcodes, the origin of the short-reads fragments can be determined, and long-range information can be inferred. Such reads thus combine the high-quality of the short-reads with the long-range information of the long-reads. As a result, Linked-Reads are particularly useful for various applications, such as genome scaffolding [4], and especially SV calling, on which we further focus below.

### 1.1 Related works

Since their inception, several methods were developed to detect SVs using Linked-Reads. These methods mainly focus on the detection of large SVs (around 10 kb) by leveraging the long-range information of the Linked-Reads. As of today, the nine following tools are available: Long Ranger [1,5], GROC-SVs [6], LinkedSV [7], NAIBR [8], VALOR/VALOR2 [9,10], ZoomX [11], Novel-X [12] and the NUI-pipeline [13]. Most of these methods rely on pairwise comparison of regions of the reference genome, in order to retrieve pairs of distant regions that share a higher number of barcodes than what would be expected based on their distance. Indeed, such region pairs indicate regions that actually appear close to each other on the resequenced genome, since adjacent regions are expected to share more barcodes than distant ones, and thus represent potential SV evidence. However, these methods do not make use of efficient barcode indexing strategies. As a result, they either need to store the barcodes of each region, which can be extremely memory consuming, or extract the barcodes from the same region multiple times, which can be highly time consuming.

Moreover, all of the aforementioned methods were mainly designed for human data, and especially, all of them were tested exclusively on human datasets in their respective publications. As a result, this focus on human data, and the lack of indexing strategies lead to scalability issues and to a poor applicability to non-model organisms, for which reference genomes are highly fragmented or sequencing data display high levels of heterozygosity. For example, Long Ranger displays an error message and cannot run on reference genomes composed of more than 1,000 contigs, while tools such as LinkedSV and VALOR2 can require up to more than 1 TB of RAM, and others such as GROC-SVs and NAIBR sometimes undergo an indefinite sleep after running for a few days.

### 1.2 Contribution

We introduce LEVIATHAN, a new Linked-Reads based SV calling method that aims to overcome these limitations, and especially mitigate resource consumption, and allow applications to non-model organisms. To achieve scalability, our method relies on a new indexing strategy, that allows to record the occurrence positions of the different barcodes through the input BAM file. This index allows to quickly and efficiently compute the number of common barcodes between all possible pairs of region that share at least one barcode. The numbers of shared barcodes between region pairs thus give a first hint as to where SVs might be located. In a second step, classical short reads methods, such as discordant paired-reads and split reads analysis are applied to region pairs sharing a sufficient number of barcodes, to further filter out false-positives, and accurately determine the types and breakpoints of actual SVs.

Our method can thus detect large SVs of at least 1,000 bp, including deletions, duplications, inversions and translocations. However, it has no support for novel insertions yet. Our experiments on simulated data show that it compares well to the state-of-the-art in terms of recall and precision, and also in terms on resource consumption. Moreover, experiments on real data also show that it manages to run on non-model organisms on which other tools either fail to run or require unreasonable amounts of resources. LEVIATHAN thus allows to analyze a wider range of datasets than the state-of-the-art, and opens doors to broader analysis of SVs in a large variety of organisms.

## 2 Methods

### 2.1 Overview

LEVIATHAN takes as input a BAM file representing the alignments of the sequencing reads of interest against a reference genome. This BAM file can either be generated by a Linked-Reads dedicated mapper, such as Long Ranger, or by any other aligner. However, when using another aligner, the reads require pre-processing prior to alignment, in order to extract the barcodes from the sequences and append them to the headers. For instance, such a pre-processing can be performed using Long Ranger basic.

LEVIATHAN relies on two distinct steps. The first step relies on the computation of the amount of common barcodes between region pairs of the reference genome, in order to highlight SVs candidates. The second step then acts as a refining step, and relies on classical short read methodologies to further filter out erroneous candidates and determine the types and breakpoints of actual SVs. An overview is given in Figure 1, and the different steps are further in the following subsections.

**Fig. 1.**
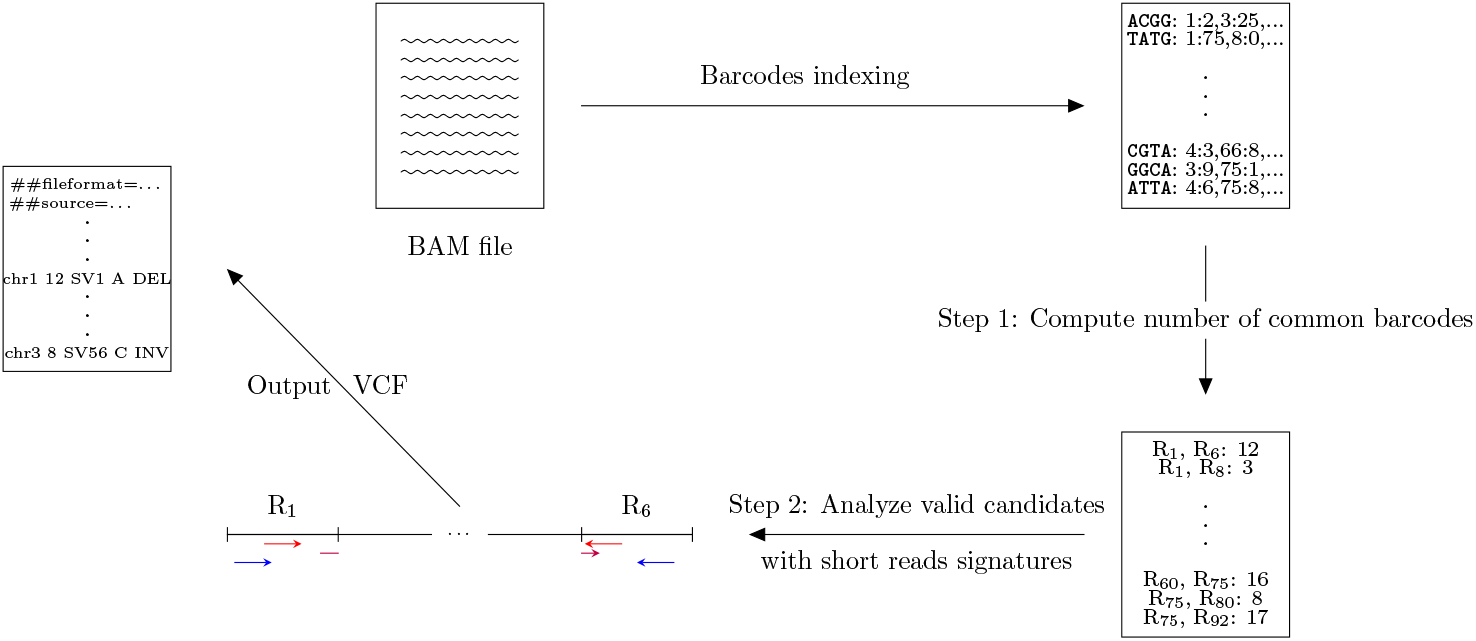
Overview of the workflow of LEVIATHAN. First, the occurrence positions of the barcodes appearing in the BAM file are indexed. The first step queries the index, to identify region pairs that share at least one barcode, and compute the number of common barcodes of such pairs. The distribution is then analyzed, and a threshold above which region pairs are further considered is defined. The second, refining step, analyzes reads signatures of these pairs, in order to define the types and the breakpoints of the SVs, which are output in VCF format.

### 2.2 Index construction

The index construction step relies on LRez [14], a tool and library designed to process Linked-Reads barcodes, which, among other functionalities, provides indexing features. LEVIATHAN thus uses LRez to build an index containing the occurrence positions of each barcode in the BAM file. This index is stored as a map, associating each barcode (in binary representation, 2 bits per nucleotide) to its list of occurrence positions in the reference genome, in format chromosome:position.

### 2.3 Computing the number of common barcodes between region pairs

Once the barcode index is built, the reference genome is divided in non-overlapping regions of size *L (L* = 1,000 by default, although this can be user-defined). As a result, LEVIATHAN only considers SVs whose breakpoints are located more than L bp apart on the reference genome. Iterating through the index then allows to easily identify region pairs that share at least one common barcode, as well as the exact number of common barcodes between such region pairs. This information is stored in a map, where the key is a region pair, and where the value is the number of common barcodes these regions share. This indexing and querying strategy allows to avoid the explicit comparison between every possible region pair, and thus allows a massive speed-up. This indexing strategy allows to address a major issue of other state-of-the-art tools, since direct comparison of region pairs’ barcodes, which either require to store the barcodes of each region or extract the barcodes of each region multiple times, and which is either time or memory consuming, is avoided.

However, processing the index as such still raises a memory issue, since numerous region pairs will share only few barcodes by chance, and will need to be stored, despite the fact they will not be considered as candidates in further steps due to their weak support. To mitigate memory consumption, we iterate through the index N times *(N* = 10 by default). Given *R* is the total number of regions in the reference genome, for each iteration, we only compute the number of common barcodes between region pairs for which the first region is comprised between the ((*i* − 1) * *R/N* + 1)-th and the (*i* * *R/N*)-th region of the reference genome. At the end of each iteration, the region pairs that share less than *B* barcodes (*B* = 1 by default) are removed, since, as previously mentioned, they will not be considered as candidates in further steps. We set this value to 1 by default, since, for Linked-Reads a given barcode does not correspond to a single molecule, but can correspond to up to 10 different ones. As a result, it is frequent that two distant regions share a given barcode by chance, but it is much less likely that they share more. Moreover, with *B* = 1, this filter allows to filter out as much as 97% of the overall number of region pairs, thus greatly reducing memory consumption. During this step, we also gather statistics on the empirical distributions of the number of shared barcodes between region pairs according to their distances, which will be used in the following step.

### 2.4 Identifying region pairs with high numbers of common barcodes

Once the numbers of common barcodes between all the pairs of regions have been computed, we need to define a threshold above which region pairs will be considered as putative SVs candidates. Effectively, despite the fact region pairs sharing only few barcodes are dynamically filtered while querying the index, a large number of pairs will still share a low number of barcodes, simply by chance in the absence of any SV. Further analyzing all of them would thus require an unreasonable amount of resources. Moreover, it is worth noting that pairs of regions that appear close to each other on the reference genome will naturally share more barcodes than pairs of regions that either appear far from each other, or even more so, on different chromosomes. As a result, considering all region pairs as one to analyze the distribution of their numbers of shared barcodes, and thus defining a single threshold, could be misleading and lead us to ignore distant pairs of regions that contain a SV breakpoint, but share an insufficient number of barcodes.

To compensate, we consider three distinct classes of distances between regions in a pair: the pairs of regions that are located close to each other (either directly adjacent or separated by on region at most), the pairs of regions that are moderately distant (separated by two to ten regions), and the pairs of distant regions (separated by more than ten regions) or on different chromosomes.

For each distance class, we then chose the 99-th percentile of the empirical class distribution as a threshold. Candidate region pairs that share less barcodes than their associated threshold are removed and not further considered.

Finally, if a region is involved in an excessively high number of pairs (> 1,000 by default), all of its pairs are also removed from the candidate list. Indeed, such regions are most probably either involved in multi-mapping problems, or prone to erroneous mapping caused by repeated regions. As a result, they are thus filtered out, since the probability of a single region being involved in such a large number of SVs is feeble. Moreover, excluding such regions from further analysis once again helps us reducing computation times. Other regions passing all these filters are then independently processed.

### 2.5 Candidate SV processing

For each of the candidates passing the previously described filters, LEVIATHAN then investigates regular short reads signals. First, reads that map on both regions are retrieved, and only these reads are then further analyzed. Classical short-reads methods are thus applied to analyze discordant paired-read and split read signatures between these two regions, in order to further determine whether a candidate is a valid SV or not, and to identify the type and the breakpoints coordinates of the actual SVs. Figure 2 illustrates the relationship between short reads signals and SVs types.

**Fig. 2.**
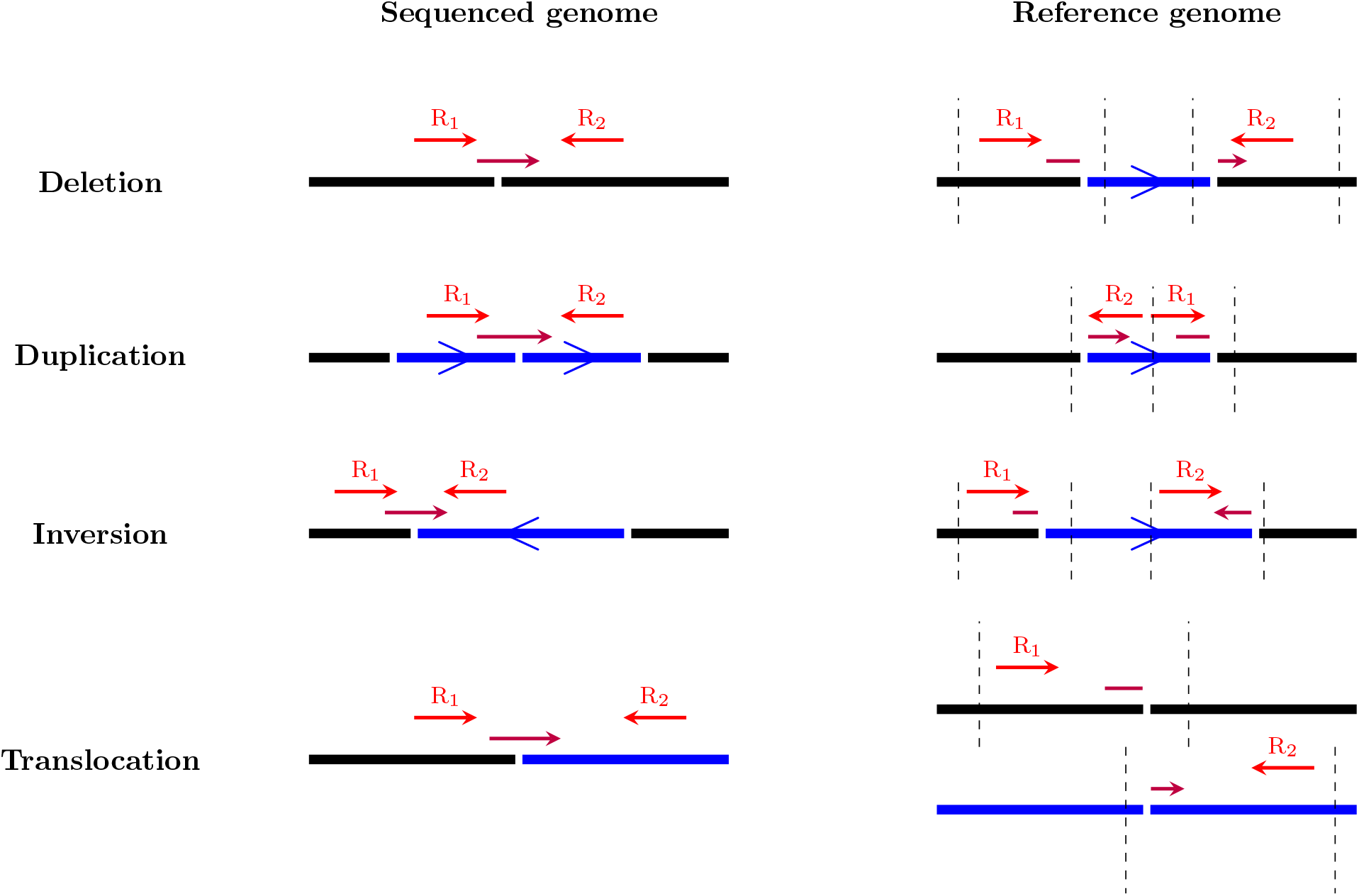
Short reads signals (discordant paired-reads and split reads) used to discriminate SV types when analyzing candidates. Dashed lines represent the regions which are considered when analyzing short-reads signals.

From this analysis, multiple support values are associated to each candidate. These values register the number of shared barcodes between the two regions, the support of each SV type, the overall number of discordant paired-reads, the overall number of split reads, as well as the support of all the possible breakpoints, in each of the two regions. A candidate is then considered as a valid SV if its support values are sufficiently high. By default, the minimum required supports are at least one discordant read pair, and at least one split read, indicating the breakpoint of the SV, in each of the two regions. Candidates passing these two filters are thus considered as valid SVs.

In terms of implementation and optimization, candidates are sorted in such a way that region pairs in which a same given region is involved are gathered together, to that the alignments of that said region only need to be extracted once. Additionally, each candidate (region_1_, region_2_) is gathered with the other candidates of the region involved in the largest number of candidates. For instance, if region1 is involved in three candidates, and region_2_ is involved in five, the candidate (region_1_, region_2_) will be gathered with other candidates of region_2_. This allows to further reduce the number of alignments extractions, and thus further optimize the runtime.

### 2.6 SV filtering and output

Prior to output, a last filtering step is applied to SV candidates. It can happen that the same SV event is represented by several breakpoint pair candidates whose genomic coordinates are very close (within 10 bp from each other) or identical, but with a different annotated type. In such a case, we only report the candidate with the largest cumulative support (barcode and short-read signal supports).

LEVIATHAN finally outputs its final list of SVs in VCF format, reporting detailed information regarding the SV, such as its type, its beginning and end positions, its length, the number of barcodes shared between the two involved regions, and the number of discordant paired-reads involved.

## 3 Results

We evaluated LEVIATHAN on simulated and real datasets. For the real data, we chose a dataset from the butterfly *Heliconius numata*, which is a non model organism and for which the discovery of structural polymorphism is of special interest. In this mimetic butterfly, it was shown that several large inversions in a 1.3 Mb locus are associated to its wing color pattern and then play a crucial role in its population biology [15]. As a non model organism, it does not have a chromosome-level reference genome, instead its 360 Mb draft genome is rather fragmented with a total of 16,950 contigs (N50: 474 kb). This genome, together with 10X genomics whole genome resequencing data of 12 individuals are available under the PRJNA676017 project ID on NCBI.

In order to properly evaluate the results quality, we also simulated data with controlled sets of SVs. To do so, we used LRSim [16] to ensure producing data that mimic the actual characteristics of Linked-Reads. We generated two datasets, one from the *H. numata* genome and one from the *H. sapiens* GrCh38 chromosome 1 (250 Mb), in order to compare the results between non-model and model organisms with similar genome sizes but different genome complexities. However, since LRSim had troubles simulating data on the highly fragmented assembly of *H. numata*, we had to filter out contigs shorter than 27,500 bp, resulting in a reference genome composed of 1,054 contigs (total size: 272 Mb and N50: 924 kb). Both datasets were simulated with a 30x coverage, and contained respectively 1,348 SVs and 1,048, ranging from 1,000 to 100,000 bp, including, deletions, duplications, and inversions as well as translocations for the *H. numata* dataset. No insertions were simulated, since LEVIATHAN is not currently able to process them. Additionally, SNPs were also inserted, in order to further mimic real data.

We compared LEVIATHAN against other state-of-the-art Linked-Reads SV calling tools. All tools were ran with default or recommended parameters, and using 8 threads. Moreover, both simulated and real data were used in our experiments.

### 3.1 Validation of the method with simulated data

To precisely asses the accuracy of the different tools, we first tested them on the simulated datasets, where precise recall and precision could be computed. However, on both datasets, we could not manage to get GROC-SVs, LinkedSV and Valor to run properly. Indeed, GROC-SVs and LinkedSV crashed on both datasets, while Valor also crashed on the *H. numata* dataset, and ran for more that two days on the *H. sapiens* dataset.

For these experiments, a SV was validated as a true positive if its breakpoints were correctly predicted, within 100 bp from a true SV reported in simulation files. Since NAIBR only reported the breakpoints of the SVs it detected, we did not take into account the SV types, in order to allow fair comparison. Statistics of the aforementioned tools on the two simulated datasets, along with their runtime and memory consumption are reported in Table 1. For the fast mode of LEVIATHAN, the 99-th percentiles of the distributions were chosen, while for the sensitive mode, the 95-th percentiles were chosen.

**Tab. 1.** Results reported by the different SV calling tools on the two simulated datasets. ^1^ GROC-SVs crashed after 6 hours. ^2^ LinkedSV crashed after 19 minutes. ^3^ Valor was killed after 2 days of computing. ^4^ LinkedSV crashed after 23 minutes. ^5^ Long Ranger could not be run on the *H. numata* dataset, since it does not allow the reference genome to contain more than 1,000 contigs. ^6^ Valor crashed upon start.

| Dataset | Tool | Recall (%) | Precision (%) | Time | Memory (MB) |
| --- | --- | --- | --- | --- | --- |
| *H. sapiens* (chr 1) | GROC-SVs<sup>1</sup> | - | - | 6 hours | 11,030 |
|  | LEVIATHAN (fast) | 71.18 | **97.14** | **11 min** | **4,603** |
|  | LEVIATHAN (sensitive) | **89.03** | 92.38 | 37 min | **4,603** |
|  | LinkedSV<sup>2</sup> | - | - | 19 min | 9,311 |
|  | Long Ranger | 72.04 | 50.23 | 11 h 29 min | 7,062 |
|  | NAIBR | 68.13 | 78.12 | 44 min | 25,764 |
|  | Valor<sup>3</sup> | - | - | > 2 days | - |
| *H. numata* | LEVIATHAN (fast) | 63.65 | **97.39** | **8 min** | **4,560** |
|  | GROC-SVs | - | - | 24 min | 1,403 |
|  | LEVIATHAN (sensitive) | **66.54** | 92.95 | 13 min | **4,560** |
|  | LinkedSV<sup>4</sup> | - | - | 23 min | 30,250 |
|  | Long Ranger<sup>5</sup> | - | - | - | - |
|  | NAIBR | 40.73 | 86.19 | 10 min | 11,736 |
|  | Valor<sup>6</sup> | - | - | - | - |

Results on the *H. sapiens* dataset show that, in terms of resource consumption, LEVIATHAN was faster than Long Ranger and NAIBR, both in fast and sensitive mode, and also required less memory, especially compared to NAIBR. However, it is worth noting that Long Ranger does not accept BAM files as an input, and that its reported runtime thus also includes reads mapping. In terms of recall, Long Ranger performed slightly better than LEVIATHAN (fast), but reached a much lower precision. Compared to NAIBR, even in fast mode, LEVIATHAN reached both higher recall and higher precision. Moreover, it is also worth nothing that, in sensitive mode, LEVIATHAN reached up to 89% of recall, while still running faster than NAIBR, and consuming the same amount of memory as in fast mode. Precision was however lower in sensitive mode than it was in fast mode, which can be explained by the fact that, in sensitive mode, a larger number of SV candidates are considered, which can increase the false-positives rate. Nonetheless, LEVIATHAN still reached more than 92% of precision in both modes, and largely outperformed both NAIBR and especially Long Ranger.

On the *H. numata* dataset, Long Ranger could not be run since it does not allow the reference genome to contain more than 1,000 contigs. Once again, the two modes of LEVIATHAN required almost three times less memory than NAIBR, and LEVIATHAN (fast) also ran faster. In terms of recall and precision, both modes of LEVIATHAN outperformed NAIBR, whose recall was particularly low, failing identifying more than half of the SVs (recall of 40.73%). In comparison, LEVIATHAN (fast) reached a recall of 63.65%, while running faster than NAIBR, and LEVIATHAN (sensitive) reached a recall of 66.54%, despite requiring a slightly larger processing time than NAIBR. In terms of precision, both modes of LEVIATHAN once again outperformed NAIBR, reaching up to 97.36% in fast mode. While LEVIATHAN still outperformed NAIBR on this dataset, its overall performance was not as good as on the *H. sapiens* dataset. Although this can be partially explained by the low quality of the reference genome we used, this still leaves us room for improvement, and future works should thus head in the direction of studying why such a proportion of SVs remained undetected.

### 3.2 Application to a real butterfly dataset

We then applied the different tools on the real dataset of *H. numata*. On this dataset, no tool except LEVIATHAN managed to run. Indeed, Long Ranger could not be run since the reference genome was composed of more than 1,000 contigs, GROC-SVs and NAIBR were stopped after 15 days of computation, Valor crashed during processing, while LinkedSV also crashed and required more than 1 TB of RAM.

In comparison, LEVIATHAN managed to run in less than two hours, only required 18 GB of RAM, and reported a total of 50,000 SVs. On this dataset, we were especially interested in finding inversions located in the *supergene* locus, the locus associated to the wing color patterns of *H. numata*. On the particular individual we studied in this experiment, LEVIATHAN did report the 430 Mb inversion at its expected breakpoints. This inversion was initially detected with SNPs, as it is associated to strong sequence divergence between individuals that display or do not display it. It was then further confirmed via PCR, and breakpoints were refined by aligning different genome assemblies [15].

While other variants reported by LEVIATHAN still need to be properly analyzed, its ability to run on such non-model organisms, for which the reference genome are highly fragmented, without requiring an unreasonable amount of resources, represents a major improvement compared to the state-of-the-art. Moreover, the fact that the inversion of interest could be detected at its expected breakpoints is particularly promising for the analysis of the remaining reported SVs.

## 4 Discussion and conclusion

We presented LEVIATHAN, a new SV calling tool for Linked-Reads data. LEVIATHAN makes use of a barcode index, which allows to quickly and efficiently compute the number of shared barcodes between pairs of regions of the reference genome. By identifying the region pairs than share a high number of barcodes, LEVIATHAN is then able to highlight pairs of regions that represent potential SVs. Complementary classical short reads methods are then applied, in order to further analyze such pairs of regions, and determine whether they are actual SVs, as well as their types and breakpoints in such cases.

Out experiments show that LEVIATHAN compares well to the state-of-the-art in terms of recall and precision, all the while being faster and consuming less memory. Moreover, LEVIATHAN also allows to analyze non-model organisms on which other tools do not manage to run or require an unreasonable amount of resources. As a result, LEVIATHAN thus tackles the main limitations of other Linked-Reads SV calling tools, and allows to better scale to large datasets, as well as to accurately analyze non-model organisms, for which assemblies are composed of large numbers of contigs. We thus believe LEVIATHAN could allow to discover new sets of SVs on a broad range of such datasets, which would be a major improvement over existing tools.

As future work, we are, first of all, planning to try LEVIATHAN on several other non-model datasets in order to try to detect new SVs. Moreover, we are also planning to integrate a local assembly feature, that would not only help us detect SV breakpoints more accurately, but that would also allow us to detect insertions. Other optimizations, such as analysis and comparison of SVs sharing common breakpoints, to better determine their types, and in-depth study of undetected SVs in simulated data are also currently being investigated. Finally, we are considering integrating other short-reads classical approaches to LEVIATHAN, in order to allow the detection of shorter SVs. Since most Linked-Reads based SV calling tools only focus on large SVs, this would allow to detect a broader range of SVs, while having to run a single tool.

## Acknowledgements

This project has received funding from the French ANR ANR-18-CE02-0019 Supergene grant. We acknowledge the GenOuest bioinformatics core facility (https://www.genouest.org) for providing the computing infrastructure. We warmly thank Mathieu Joron and Paul Jay for sharing their data and results on Heliconius inversions.

